# Distributed context-dependent choice information in mouse dorsal-parietal cortex

**DOI:** 10.1101/2021.03.02.433657

**Authors:** Javier G. Orlandi, Mohammad Abdolrahmani, Ryo Aoki, Dmitry R. Lyamzin, Andrea Benucci

**Affiliations:** RIKEN Center for Brain Science, 2-1 Hirosawa, Wako-shi, Saitama 351-0198, Japan; University of Tokyo, Graduate School of Information Science and Technology, Department of Mathematical Informatics, 1-1-1 Yayoi, Bunkyo City, Tokyo 113-0032, Japan

## Abstract

Choice information appears in the brain as distributed signals with top-down and bottom-up components that together support decision-making computations. In sensory and associative cortical regions, the presence of choice signals, their strength, and area specificity are known to be elusive and changeable, limiting a cohesive understanding of their computational significance. In this study, examining the mesoscale activity in mouse posterior cortex during a complex visual discrimination task, we found that broadly distributed choice signals defined a decision variable in a low-dimensional embedding space of multi-area activations, particularly along the ventral visual stream. The subspace they defined was near-orthogonal to concurrently represented sensory and motor-related activations, and it was modulated by task difficulty and contextually by the animals’ attention state. To mechanistically relate choice representations to decision-making computations, we trained recurrent neural networks with the animals’ choices and found an equivalent decision variable whose context-dependent dynamics agreed with that of the neural data. In conclusion, our results demonstrated an independent decision variable broadly represented in the posterior cortex, controlled by task features and cognitive demands. Its dynamics reflected decision computations, possibly linked to context-dependent feedback signals used for probabilistic-inference computations in variable animal-environment interactions.

## Introduction

Choice signals in the brain reflect a relationship between neural activity and the animal’s choice during decision making [1]. Previous research has focused particularly on perceptual behaviors, using “choice probability” as a metric to quantify correlations between the activity of neurons and the trial-to-trial fluctuations in animals’ choices [2]. The inferred correlations were usually computed by factoring-out task regressors that might inform choices; for instance, a frequently used paradigm is the random-dot discrimination task, used to record from neurons in middle temporal visual area (MT) [3]. Since in this area neurons clearly encode the direction of stimulus motion, choice probability is measured at near-zero coherence.

Choice signals have also been described using a decision variable (DV) derived from neural responses that dynamically followed the decision state of the animal [4]. In evidence accumulation tasks—as the random dots task—the DV is driven by sensory evidence as well as by the animal’s “internal state”, with weak sensory evidence best revealing internally driven computations, such as changes of mind [5]. In posterior cortical regions, choice signals have been characterized by a large degree of heterogeneity [6–10]. Most studies have focused on individual regions, and found that numerous variables can influence the probability of detecting these signals; for instance, the stimulus-coding strength of neurons [6, 11], the correlation properties of the network [12, 13], the area location in the sensory hierarchy [14, 15] and even the strategy used by the animal to solve the task [16]. Furthermore, choice signals can be difficult to disentangle from coincident task- and behavior-related processes, as those associated with the execution of actions, or with modulatory signals reflecting variability in the attention state of the animal [17]. This latter process can contextually enable, route, and gate choice information [18], thus greatly affecting how choice signals are distributed in these areas. It is therefore possible that during a task choice signals are simultaneously represented across multiple areas, with amplitudes that, relative to concurrently represented processes, are areas-specific and contextually modulated by the internal state of the animal [19].

In this work, we selected a task, recording methodology, and analytical framework that enabled us to examine this possibility. As the animals engaged in a complex variant of a two-alternative forced choice (2AFC) orientation discrimination task [20], we isolated choice signals from sensory and motor-related activations by performing mesoscale imaging of neural responses (GCaMP) from the mouse posterior cortex. Using recent tensor decomposition methods [21] and activity-mode analysis [22], we isolated choice signals from concurrently represented sensory and motor-related activations, and characterized their distinct spatial and temporal properties across cortical areas. In a reduced space of multi-area activations, choice signals defined an embedding subspace for left or right (L/R) decisions that was near-orthogonal to that of sensory signals and movement components and was modulated by task difficulty. In visual areas, their spatial signature was prominent in ventral stream regions, the areas implicated in the processing of stimulus’ identity. When monitoring fluctuations in the attentional state of the animals, we found that sustained attention differentially modulated the choice signal, but the embedding subspace remained invariant. Modeling the animal behavior with recurrent neural networks (RNNs [23–25]), trained with the animals’ trial-by-trial decisions, provided mechanistic evidence that the context-dependent representational dynamics reflected the computations underlying the 2AFC task.

## Results

### Mesoscale imaging of dorsal-parietal areas during a discrimination task

Using an automated setup featuring voluntary fixation of the animals’ heads [26] (Fig. 1a), we trained mice (*n* = 7) to carry out a complex version of a 2AFC orientation discrimination task [20]. The animals had to use their front paws to rotate a toy wheel [27] that controlled the horizontal position of two circular grating stimuli presented on a screen positioned in front of them. Each stimulus was presented at monocular eccentricities with orientations that varied from trial to trial. To obtain a water reward, mice had to shift the stimulus most similar to a learned target orientation to the center of the screen (Fig. 1b, c), with the actual learned orientation rarely shown to the animal. Therefore, difficulty had an invariance to absolute orientations, which had to be ignored by the animal, and depended only on the relative orientation between the two stimuli. After reaching performance levels above 75 % correct (Fig. 1d), we used a macroscope to image mesoscale GCaMP responses in approximately 12–14 dorsal-parietal cortical areas (Fig. 1e; Methods). In individual trials, the neural activity was highly variable, with response activity associated with the onset of visual stimuli and movements of the limbs, trunk, and eyes, as recently described [28] (Fig. 1f).

**Fig. 1.**
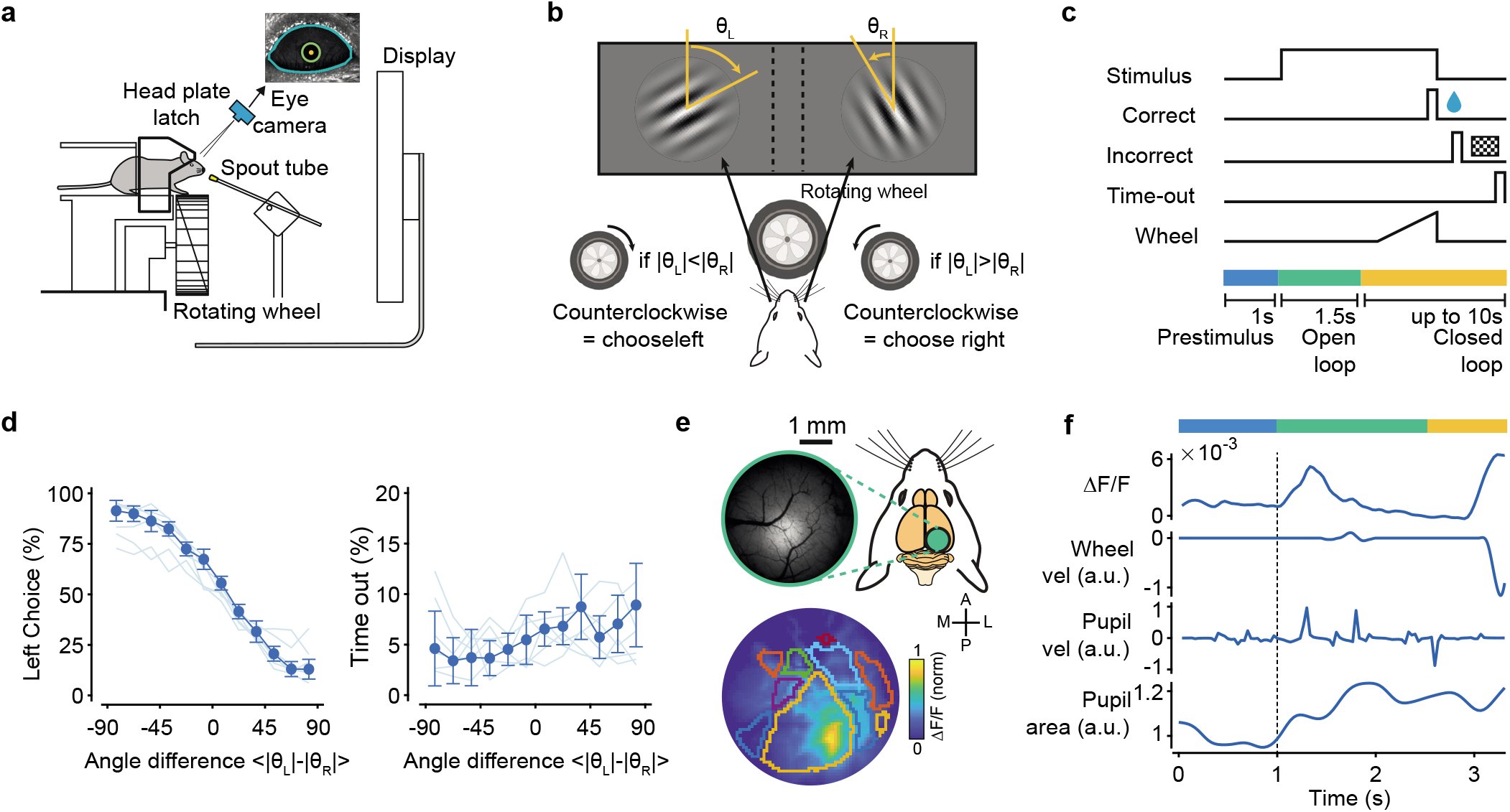
Imaging dorsal-parietal areas during an orientation discrimination task. **a**, Mice were trained on a two-alternative forced choice orientation discrimination task (2AFC) using an automated setup featuring voluntary head fixation. They signaled a L/R choice by rotating a toy wheel with their front paws. **b**, Mice rotated the wheel to position the most vertical of two oriented gratings in the center of the screen. **c**, Trial structure: after a 1s pre-stimulus period the stimulus was presented, followed by a 1.5 s of open loop (OL) interval where wheel movements were decoupled from stimulus movements. Afterwards, in the closed loop period (CL), wheel rotations resulted in L/R horizontal shifts of the stimuli. Correct choices were water rewarded; Incorrect choices were followed by a checkerboard pattern presentation. 10 s of no movements in the CL triggered a time out). **d**, Left: mice performance in the task (fraction of left choices) as a function of angle difference from the target orientation (nominal value of zero). Thick line, mean (± s.e.) across animals; thin lines, individual animals. Right: fraction of timeout trials as a function of difficulty. Timeout trials did not depend on task difficulty, **e**, Widefield calcium imaging of the posterior cortex of Thy1-GCaMP6f mice, with retinotopic mapping of 10-12 visual areas (colored contours). **f**, Simultaneously recorded fluorescence signals (Δ*F/F*), wheel and eye velocities and pupil area. In this example, choice was signaled at t = 3.1s (sharp increase in wheel velocity).

### Decomposition of neural responses

To extract different variables from the neural signal and map them to defined cortical regions, we adopted a recent variant of non-negative matrix factorization—locaNMF [21]. This decomposition method identifies tensor components associated with specified seeding regions. When seeding on a given area, locaNMF decomposes the signal into a sum of separable spatial-temporal tensors, with spatial components constrained by the seeding region and temporal components representing the scaling amplitudes of the spatial components. These temporal vectors are potentially more informative than a single vector computed as the average across spatial locations (pixels) within a given area [21]. We aligned all imaging sessions according to the Allen Common Coordinates Framework (CCF [29], Fig. 2a) and seeded the initial spatial decomposition using 10 large regions centered on retinotopically identified areas that extended significantly beyond area boundaries (Extended Data Fig. 1). Consistent with the initial seeding, the factorization typically converged toward components with peak amplitudes within individual retinotopic areas (Fig. 2b). Depending on the seeding region, associated temporal components differentially emphasized sensory or behavioral variables; for instance, when seeding on the primary visual cortex, the largest component (in explained variance, EV) clearly highlighted a stimulus-evoked response (Fig. 2b). However, the largest components within parietal regions [30] (e.g., A, RL) showed negligible visually driven responses and strong movement-related activations (Fig. 2b). Each locaNMF component provided significant explanatory power, with each main component of a seeding area contributing, on average, 9.6% of the total EV (Extended Data Fig. 1). By contrast, the first PCA component contributed, on average, approximately to 85 % (Extended Data Fig. 1), being strongly influenced by large amplitude movement-related activations [31]. For each area, the proportionality between the surface area and the number of components significantly contributing to the EV (Methods) was not always straightforward; for instance, areas AL and L had commensurate surface area and contributed similarly to the overall EV, but L required about twice as many components as AL (Extended Data Fig. 1). To identify task- and behavior-related variables in locaNMF components, we defined state vectors in a multi-dimensional space of component activations (Fig. 2c). This approach further reduced the dimensionality of the data by isolating activity dimensions that linearly discriminated pairs of variables. To examine components associated with visual signals, we defined a stimulus axis as the difference between vectorized tensor components in the presence and absence of the stimulus (Methods). This axis remained stable after the stimulus’ appearance (Extended Data Fig. 2) and the projected locaNMF components deviated from the baseline about 200 ms after stimulus onset (Fig. 2d). We quantified the time-dependent increase of detectability of stimulus components using a *d*′ discriminability measure, which can be linked to Fisher information [32, 33], that bounds the variance for estimating a population-encoded parameter. Different areas contributed to *d*′ with different weights, reaching values greater than one at the peak of stimulus response (Fig. 2d, *d*′ = 1.38 ± 0.13, mean ± s.e.). Using only the LocaNMF components from a particular seeding region allowed us to also quantify the relative contribution of that area to the *d*′ discriminability. For the stimulus variable, the primary and secondary visual cortices (V1, L) had the largest discriminability (*d*′ = 1.10 ± 0.09 and *d*′ = 1.12 ± 0.13, respectively), followed by area AL (*d*′ = 0.51 ± 0.06). When attempting to discriminate the orientation of the contra-lateral visual stimulus, no area carried sufficient information, even for the most dissimilar orientation pairs (Extended Data Fig. 3), as expected from the lack of orientation domains in mouse visual cortex [34] and the mesoscale spatial resolution of our imaging system.

**Fig. 2.**
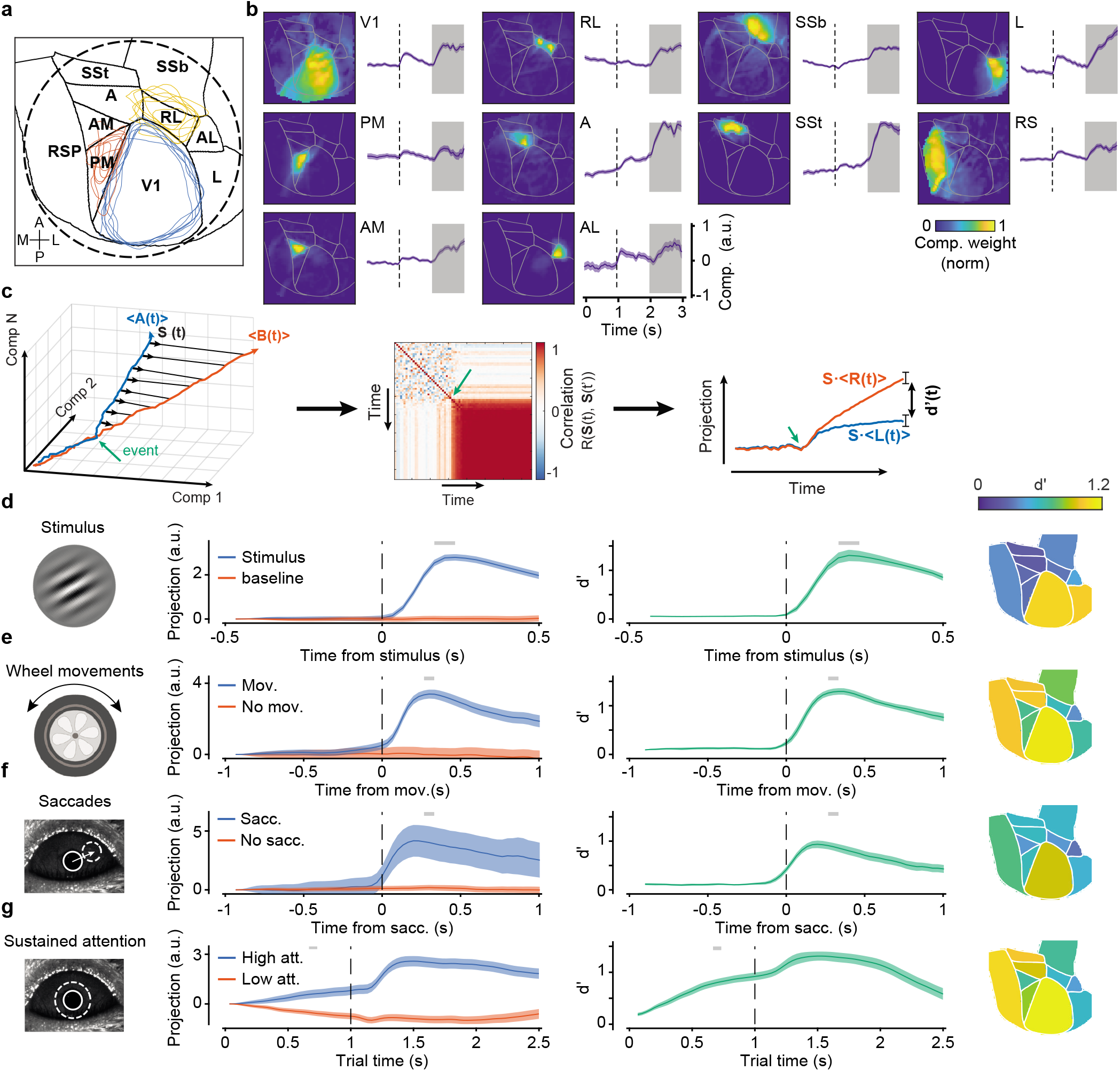
LocaNMF decomposition identifies sensory, behavioral, and attention-related variables. **a**, Characteristic imaging window (dashed circle) superimposed on 10 cortical areas from the Allen Brain Atlas reference framework. Blue, red, and yellow contours are the reference-aligned area boundaries for V1, PM and RL for each animal. **b**, Spatial weights and trial-averaged time-series of the largest locaNMF components for each of the 10 seeding regions (Extended Data Fig. 1) for a representative animal. Selected trials for the trial-average all presented wheel movements within the 1 s shaded region. **c**, Schematic for the definition of state vectors. For any given pair of task-related variables, **A**(*t*) and **B**(*t*) (i.e. locaNMF temporal components), we defined at each time point a state vector **S**(*t*) as a trial-averaged unit difference vector **S**(*t*) ~ 〈**A**(*t*)〉 – 〈**B**(*t*)〉. In this schematic, the direction of the state vector becomes stable after an event indicated by the green arrow. Vector stability is measured as the temporal autocorrelation *R*(**S**(*t*), **S**(*t*′)), (right panel). Projections (cross-validated) of the two variables **A**(*t*) and **B**(*t*) onto **S** separate over time, as quantified by a *d*′ discriminability measure. **d**, Stimulus-related state vector. Left: projections of trials with and without a stimulus response onto the stimulus state vector for a given animal. Pre-stimulus times were used for no-stimulus conditions. Lines and shaded regions indicate projection averages across trials and s.e. Middle: Discriminability *d*′ over time, averaged across all animals. Grey bars on top, epoch used for the time-average of the state vector. Right: area-specific peak *d*′ scores obtained by defining the state vector using only the components originating from that area. **e**, As in **d** for the wheel movement state vector, aligned to movement-detection time, i.e., separability between trials with and without a detected wheel movement. **f**, As in **e** for saccadic eye movements. **g**, as in **f** for sustained attention. High and low attention trials were defined based on pupil area change, using the highest or lowest 33-percentile of the area-change distribution.

Besides bottom-up visual inputs, imaged dorsal-parietal regions reflected activations associated with general movements of the body and eyes [35]. Therefore, we defined state axes associated with wheel and eye movements. Projections onto these axes resulted in high discriminability of both types of movements (Fig. 2e, f; peak *d*′ = 1.29 ± 0.07 and *d*′ = 0.94 ± 0.08 for wheel and eye movements, respectively). Area-specific projections highlighted larger contributions by anterior-medial areas (Fig. 2e, f; Extended Data Fig. 2), with *d*′ values increasing before or coincidentally with the detection of movements, suggesting pre-motor contributions (e.g., corollary discharges [36]), and reaching values greater than one after movement execution.

We also identified aspects of the variability in locaNMF components that depended on the attention state of the animal. Underlying changes in sustained attention can be both task-related (e.g., engagement or motivational state) and task independent components (e.g., arousal or alertness) [37, 38]. Accordingly, in individual sessions we observed fluctuations in performance that correlated with changes in pupil dilation and reaction times (Extended Data Fig. 4), two biomarkers associated with changes in sustained attention [39]. Based on the variability in pupil diameter (Methods), we defined a state axis that discriminated between high- and low-attention states (Fig. 2g). Associated *d*′ values deviated significantly from zero largely before stimulus onset (after imposed zero discriminability at trial onset; see Methods). Discriminability values reached *d*′ = 0.5 approximately 0.5 s after trial onset and remained above this value throughout the trial duration, with peak *d*′ = 1.31 ± 0.09. The state axis defined by attentional modulations remained stable throughout the duration of the trial (Extended Data Fig. 2), consistent with periods of high and low attention that persisted across trials [35]. The anterior-medial visual areas and the retrosplenial cortex contributed most significantly to large *d*′ discriminability (Fig. 2g, Extended Data Fig. 2).

Together, these results showed that sensory inputs, movement-related activations, and attentional signals were concurrently present in the dorsal-parietal regions, and could be separated by the locaNMF tensor decomposition, permitting the identification of their characteristic spatial and temporal signatures.

### Choice signals

This approach also allowed us to identify choice-related signals. We adopted a broad operational definition of “choice” as signals that correlated with animal’s L/R decisions, independently of the stimulus and with premotor signatures reflecting action-selection [9]. We considered trials in which the first detected wheel rotation occurred at least half a second after stimulus onset. The first detected movement after stimulus presentation did not always coincide—by definition—with the movement signaling the animal’s choice, which occurred in the closed-loop (CL, Methods) period and terminated the trial. However, we confirmed that the direction of the first movement had a large and significant correlation with the final choice ((85 ± 4) % agreement with movement directions), suggesting that the decision was made quickly after the stimulus presentation (Extended Data Figs. 4–5). We then aligned responses relative to movement times and defined a state axis that linearly discriminated clockwise from counterclockwise wheel rotations (hereafter left and right choice, respectively). LocaNMF projections onto this axis sharply separated left from right choices (Fig. 3a), reaching peak separation values approximately 0.15 s after movement detection (Fig. 3b, peak *d*′ = 1.5 ± 0.1), and with the choice axis showing two clearly stable regions, before and after movement onset (Fig. 3c). Global and area-specific *d*′ values started to increase before movement onset (Fig. 3d). We characterized pre-movement components using a piecewise linear regression analysis (Fig. 3e) applied to *d*′ curves to quantify the slope of the fit before the movement as well as the time of the slope change (Fig. 3f-g; slope = 0.19 ± 0.06 *d*′/*s*, time of slope change = (−0.06 ± 0.02) s). We found a consistent trend for positive pre-movement slopes (ramping) and pre-movement slope change times, providing evidence for distributed premovement choice components across these regions.

**Fig. 3.**
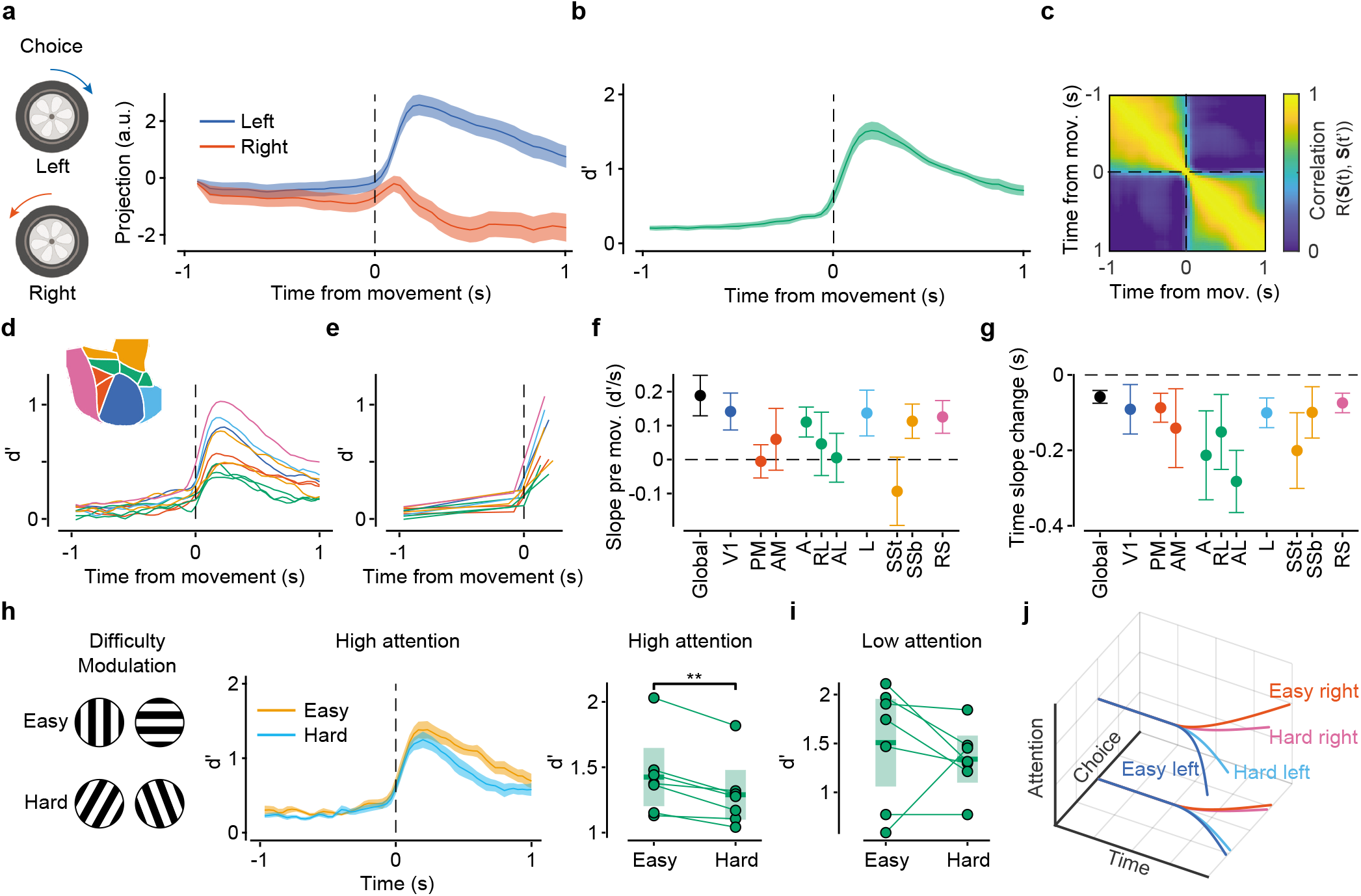
Choice signals have pre-motor component and are modulated by task difficulty and attentions. **a**, Projections on the state vector for choice signals. Wheel movements signaling either a left or a right choice were aligned to the wheel movement onset. **b**, Evolution of *d* discriminability relative to movement time. Early and late choice periods were defined relative to movement onset. **c**, Temporal stability of the state vector for choice. There is a clear change in the contribution of the choice state vector near the time of movement onset. **d**, Temporal evolution of area-specific *d* curves (inset: area color code). **e**, Piecewise linear fits of the curves in d in pre- and post-movement periods. **f**, Pre-movement slopes fitted in e for different areas; error bars, 95 % confidence intervals across animals (CI). **g**, Times of slope change for different areas from the fits in e. Data and colors as in **f**. **h**, Left: evolution of choice discriminability, *d*′, in high attention states for easy and hard trials (angle difference > or < 45°). Right: paired comparisons of peak *d*′ values from **h** for each animal (*p* = 0.003). **i**, Same as in **h** for low attention states. Paired differences were not significant (*p* = 0.50). **j**, Schematic representation of the temporal evolution of left and right trajectories with difficulty and attention

We reasoned that although evidence accumulation might not be a significant factor in our task, a decision variable [40]—reflected in the time-varying *d*′ values—would still retain its sensitivity to task difficulty. Indeed, we found that in high-attention states, *d*′ curves reflected stronger choice separation in easy trials than in difficult trials (Fig. 3h; peak *d*′ = 1.4 ± 0.1 and 1.3 ± 0.1, respectively; paired t-test, p = 0.003). In low-attention states, there was a similar trend, but the difference was not significant (paired t-test, p = 0.4). The dependence of *d*′ values on task difficulty in high-attention states revealed a modulatory effect of attention on choice signals. However, when we analyzed the average separation between trajectories across attention states, we found no significant difference (difference in peak values, paired t-test, *p* = 0.5). Furthermore, choice axes independently defined in low- and high-attention states were highly correlated (Pearson’s *r* = 0.72 ± 0.03). Together, these results indicated that the choice subspace defined by left and right trajectories remained the same across attention states, but attention enabled difficulty-dependent modulations of the trajectories (Fig. 3j).

### Distinct spatial and temporal characteristics of choice signals

Choice signals were broadly distributed across multiple areas, but with distinct spatial and temporal characteristics. We defined a spatial-distribution index (SDI) that captured whether several or only a few areas contributed prominently to the *d*′ discriminability and found that choice had the largest SDI values ((30 ± 4) %) compared to sensory, movement and attentional signals (around 10%) (Fig. 4a). Distinct interarea signatures of choice signals were also evident in the pairwise angular separation between state axes (Fig. 4b). We found that, overall, angular distances were greater than 50°, with the choice axis being near-orthogonal to the sensory, movement, and attention-related axis. In time, choice axes computed separately before (−0.1s) and after (0.3 s) movement onset were stable in the pre- and post-movement periods and orthogonal to each other. Sensory and movement components had large angles (around 70°), and the smallest angles (still over 45°) were observed between the movement and attentional axes.

**Fig. 4.**
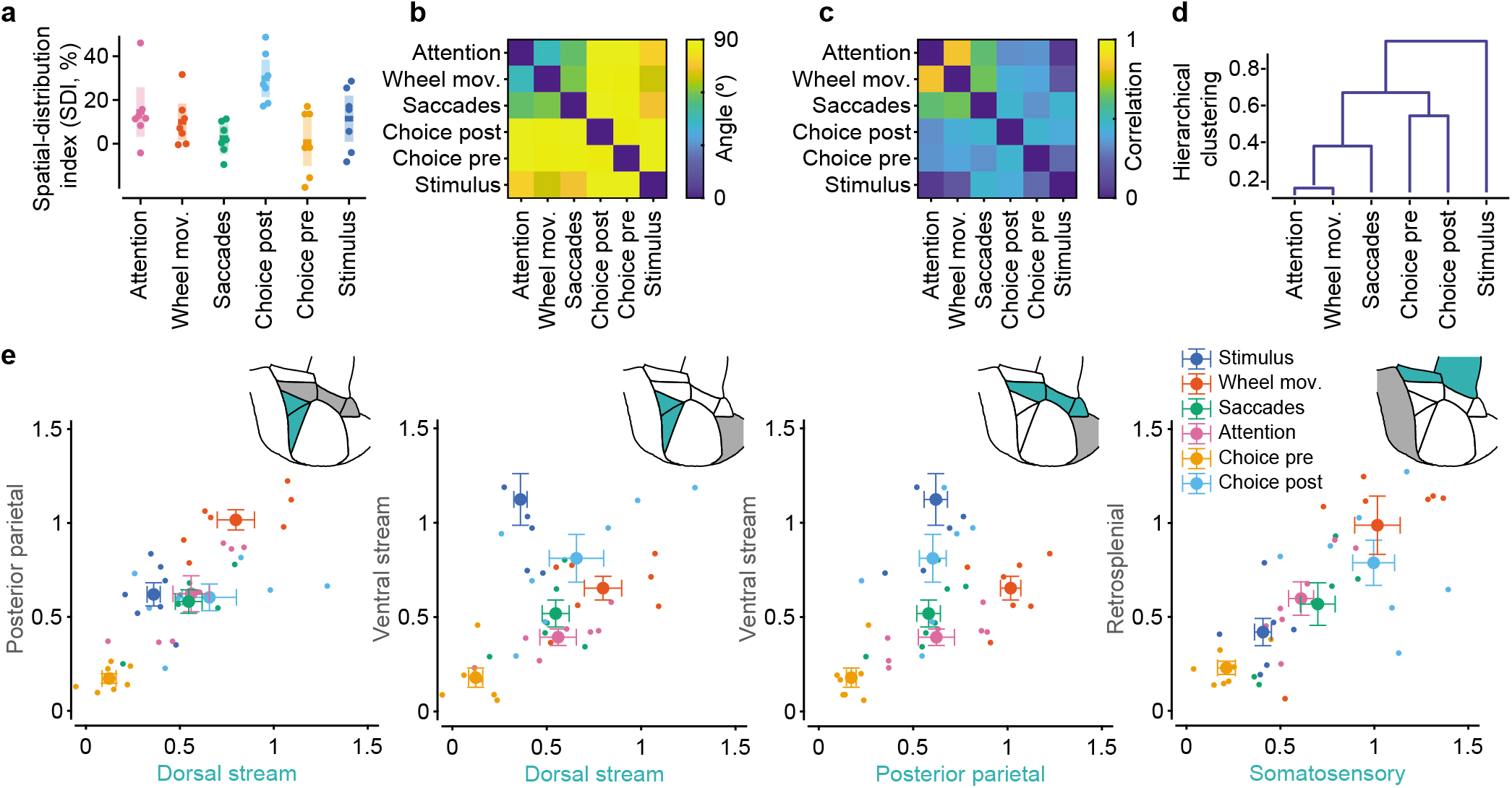
Choice is near orthogonal to other components with a ventral-stream dominance. **a**, Spatial-Distribution index (SDI) for each state vector. Choice had the largest SDI ((30 ± 4) %); dots are different animals; middle lines and shaded areas, mean and 95% CI. b, Angles between state vectors. Choice axes (pre- and post-movement) were orthogonal to all other axes (smallest angle (84 ± 7)°). Attention and wheel had the smallest angular separation (44 ±3°), followed by wheel and saccades ((56 ± 4)°). **c**, Pairwise correlations between the *d*′ indices obtained from each area used for hierarchical clustering. **d**, Hierarchical clustering from the pairwise correlations shown in **c**. Attention and wheel movements were most similar, followed by saccades. Choice pre- and post-movement onset clustered together, whereas stimulus showed the most dissimilar pattern. **e**, We computed five *d*’ values, each derived by restricting locaNMF components to one of the five area groups, thus defining a 5-D space of *d*’ components. The five broad area groups consisted of the dorsal stream (PM and AM), ventral stream (L), posterior parietal (A, AL and RL), somatosensory (SSt and SSb) and retrosplenial (RS) regions.

To examine area contributions to these differences, we performed a correlation analysis between *d*′ maps; for example, *d*′ maps for stimulus and movement components (Fig. 2e, f) had overall similar discriminability, but highlighted distinct area-specific contributions and, accordingly, the correlation between the maps was small (Fig. 4c). We systematically analyzed all pairwise correlations and used hierarchical clustering to identify components with stronger similarity in *d*′ spatial distributions (Fig. 4d). The stimulus axis was most dissimilar from others, followed by the cluster of choice axes. Attention and movement components clustered together, consistent with their individual large pairwise correlation values (*r* > 0.5, Fig. 4c).

To further examine the area-specific contributions to choice signals, we divided higher visual areas into three main groups—ventral (L), dorsal (PM, AM), and parietal (A, RL, AL) [30]—and separately analyzed somatosensory (SSt, SSb) and retrosplenial (RS) regions. V1 contributed with an overall uniform *d*′ value to all separations (Extended Data Figs. 2 and 6), hence, we did not include it in this analysis of relative differences. We then computed *d*′ values using only the locaNMF components that originated from these grouped areas and did this for all variables: visual, movement, choice, and attention. This resulted in a five-dimensional (5D: ventral, dorsal, parietal, somatosensory and retrosplenial) space, where the coordinates of a variable reflected the contribution of a given region to the *d*′ separability of that variable. When examining discriminability power in 2-D projections of this 5-D space (Fig. 4e), we found that regions contributing to choice were distinct from those of other components—including movement—being most prominent in ventral stream regions, as reflected by the large *d*′ values observed in these areas (Extended Data Fig. 6). Ventral regions significantly contributed to visual components, as expected, while dorsal and parietal regions contributed to movement variables—especially wheel movements—in agreement with a previous report [35]. Retrosplenial and somatosensory areas contributed similarly to choice and movement, with *d*′ values generally correlated across all variables (*r* = 0.8; 95 % confidence intervals (0.61, 0.97), for *d*′ correlations between somatosensory and retrosplenial areas across animals).

In summary, distributed choice signals were distinct from sensory, movement, and attentional components, dominantly in ventral-stream visual areas and modulated by task difficulty and attention, suggesting that they might reflect the decision-making computations associated with the discrimination task.

### Recurrent Neural Network model of decision dynamics

To examine this possibility, we used RNNs as implementation-level, mechanistic models of the decisionmaking process. Building on previous work showing that RNNs can capture decision-making computations associated with 2AFC discrimination tasks [23, 24], we examined the dynamics of RNNs trained according to the invariance for absolute orientations present into our task—and learned by the animals. Furthermore, we trained RNNs with the actual trial-to-trial choices of the animals, rather than using the optimal task solution, and introduced variability in attention states (Fig. 5a). Using the animals’ choices rather than the task rule, created numerous “contradictory” examples, where the input evidence for a left or right choice was non-deterministically associated with left or right output decisions, even in the easiest trials (e.g., non-zero lapse rate). As a result, although RNNs were trained to produce L/R binary choices, they learned analogue outputs that followed a psychometric probability function (Fig. 5a). Furthermore, output amplitudes depended only on task difficulty, reflecting a learned invariance for absolute orientations. Context-dependent attention modulations (introduced as an additional binary input) modified output probabilities and created shallower or steeper psychometric curves in low or high attention states respectively (Fig. 5a, Extended Data Fig. 4). Although the model was trained only with a subset of 13 difficulties and 2 attention states, it was able to generalize to any difficulty level and range of attention within the trained boundaries (Fig. 5b). We then analyzed the internal dynamics of the network by computing choice and attention axes from RNNs unit responses, as we did for the neural data with locaNMF components. In the RNNs also, the choice axis identified a decision variable that represented L/R choices as separate trajectories in a low-dimensional embedding space. Furthermore, the separation between trajectories was modulated by task difficulty, with larger separations in easy trials. This separation did not depend on absolute orientations, as expected from the RNN having learned this invariance. The choice axis was stable across levels of difficulty (Fig. 5c). Attentional modulations maintained an invariant representational geometry of the DV across the embedding space (Fig. 5d-h). This was consistent with observations of the neural data, where choice and attention axes were near-orthogonal with each other (Fig. 4b). In summary, the representational similarity between the RNN and neural dynamics, together with the lack of neural information provided to the network, indicated that the contextually-modulated choice signals observed in locaNMF components indeed represented the decision-making computations underlying the orientation discrimination task.

**Fig. 5.**
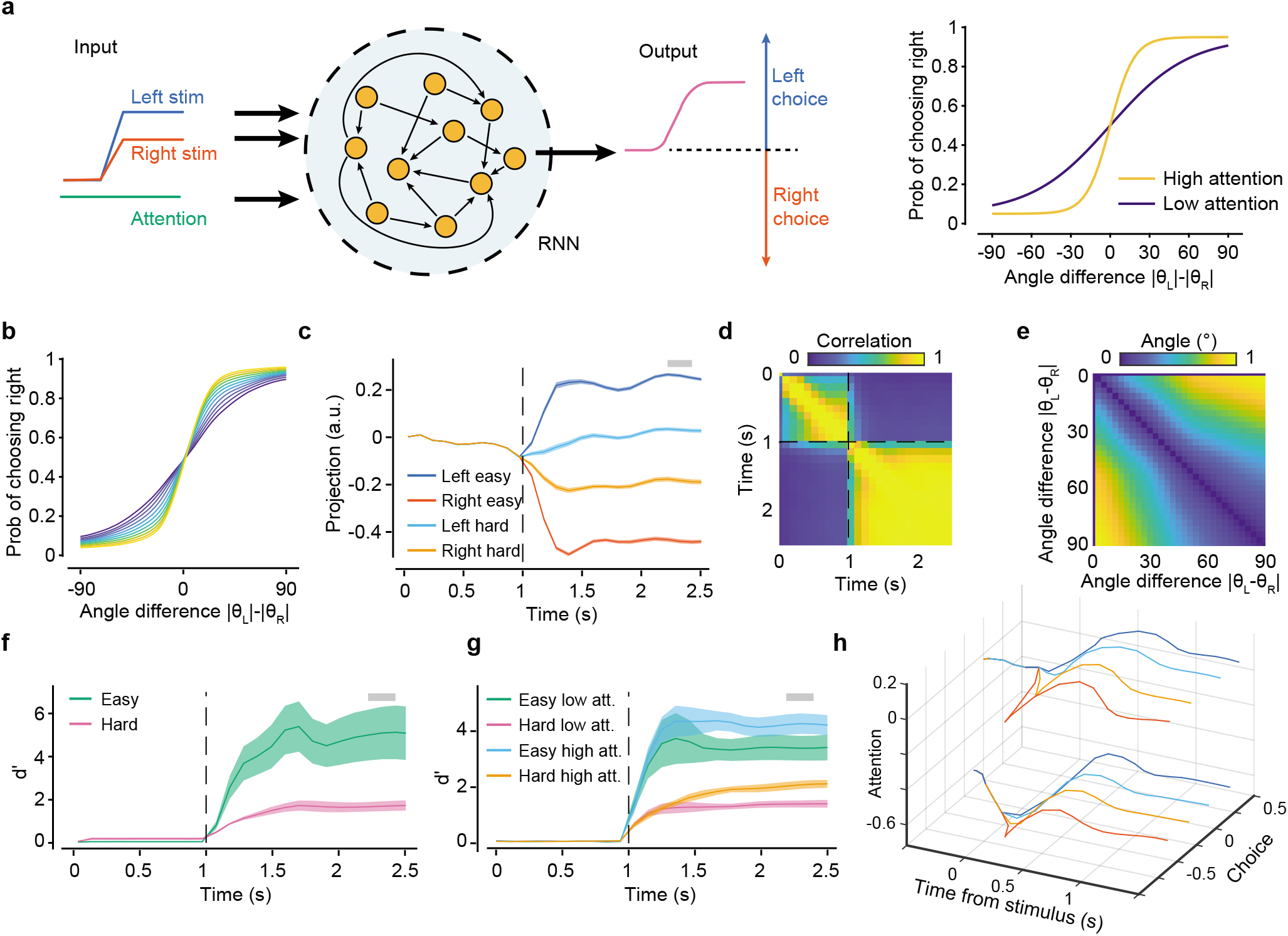
RNN model relates neural representations to DM computations. **a**, Left: recurrent neural network (RNN) architecture consisting of a module with *N* = 128 recurrently connected units. The module receives two inputs for the left and right stimuli, and one input for the attentional state. It generates a continuous output that will determine the choice. Right: Psychometric curves used to determine the proportion of L/R-choice trials in the training set for each difficulty level, depending on the attention state. **b**, Psychometric curves from the trained model showing the model generalizes to intermediate values of attention and difficulties. **c**, Projection of L/R easy and hard trials on the choice axis (state vector) following the same methods used for Fig. 2. Shaded bar denotes the selected time epoch used for state vector averaging. d, The choice axis became stable quickly after the stimulus presentation (t = 1 s). **e**, State vectors for choice, computed separately in each trial based on difficulty level, were almost parallel to each other, with the largest deviation staying below 20°. Thus, the network learned a unique choice representation that was independent of task difficulty. **f**, Discriminability for choice (*d*′) was higher in easy than in hard trials. Line and shaded area, mean and 95 % CI across 10 trained networks with different random initializations. **g**, Choice was modulated by attention, as shown by the increase in discriminability values in trials with high attention levels. **h**, Diagram of the projected trajectories in the space spanned by the axes of choice and attention.

## Discussion

Using a tensor decomposition method that retained the spatial information in the responses, and applied to mesoscale recordings in mouse dorsal-parietal cortices, we isolated choice signals that were near-orthogonal to sensory-, movement- and attention-related variables. We showed that their representational dynamics was consistent with the decision-making computations underlying the behavioral task. We also showed that choice signals, although broadly distributed, were prominent in ventral-stream visual areas and were modulated by task difficulty, with this modulation enabled contextually by the attention state of the animal. Together, these results suggest a multiplexed representation of variables in the posterior cortex, with a widespread distribution of decisional information, possibly reflecting probabilistic inference computations; for instance, information about the ongoing decision-making process could be used for perceptual inference with unreliable sensory stimuli, and to influence sensory-to-decision signal transformations that inform future action plans.

### Methodological relevance

We achieved these results by combining two powerful methods for the analysis of population responses: locaNMF and activity-mode analysis. LocaNMF reduced the dimensionality of the neural data while retaining spatial information, which would have been lost with traditional dimensionality reduction methods (e.g., PCA). Furthermore, the state space representation allowed further reduction of dimensionality while aligning the dynamics along task and decision-relevant dimensions. This latter step took place within an interpretable linear framework, where the angle between the state axes and *d*′ values could be directly linked to the linear discriminability of the underlying variables. The imaging methodology and task design used in this study facilitated the identification of distributed choice signals encoded by sparse populations of cells [41] thanks to the local integration of these responses. Furthermore, the complexity of the decision-making task might have influenced the wide spatial distribution of the representations [42].

### Feedback origin of choice signal

Choice signals emerged after stimulus onset, were broadly distributed in the posterior cortex, and could be significantly detected as early in the visual hierarchy as in V1, suggesting feedback activations from areas causally involved in the decision-making process. Other non-sensory signals identified in our recordings, e.g. related to body and eye movements, could also result from feedback activations. Indeed, feedback signals to the posterior cortex have been extensively documented in the literature, in association with a great diversity of underlying variables and computations, including attentional modulations [43], movement-associated responses [28], sensory context [44], and predictive coding [45].

### Choice signals have distinctive spatial and temporal signatures

The properties of the choice signals met several criteria that are characteristic of a decision variable. Their pre-movement components suggested that they did not simply reflect the execution of a motor plan, nor an “unsigned” pre-motor, preparatory state [46]. Choice signals did not simply reflect bottom-up, stimulus-related information that correlated with the decision process because, given the task design, the contra-lateral stimulus orientation was uninformative for L/R decisions [20], and furthermore, at the mesoscale resolution used in this study, orientations were not decodable from the neural signal [47].

Choice signals could be separated from movements. The cortical localization of movement components was prominent in dorsal-stream regions, consistent with previous reports [28, 35]. Choice signals were localized in retrosplenial cortex, and in the visual cortex, mostly in ventral stream regions, along the so-called “what” visual pathway [48]. This was consistent with the task requirements: mice had to evaluate the orientation “content” of both stimuli and make relative orientation comparisons. Absolute orientations were informative, as were the locations of the stimuli, which were the same across trials; thus, in contrast to “where” type of information, which is supposedly associated with dorsal stream regions, solving the task relied on “what” information in the ventral stream areas.

We also controlled ventral-stream responses did not link to eye movements (Extended Data Fig. 7), which typically followed whole body movements [35]. However, it is still possible that signals detected in ventral areas were associated with motor-related components that also carried choice-relevant information [49].

Attention-mediated modulations were orthogonal to the subspace defined by choice variables, with the choice axis remaining significantly autocorrelated across time irrespective of attentional state. This can be described as an isomorphic transformation in the embedding space of the decision variable (DV), where the subspace defined by the L/R trajectories is shifted, without deforming the representational geometry. The dependence of the decision variable on task difficulty was clearer in high-attention trials, but not significant in low-attention states. This might reflect an actual dependence of the decision-making process on attention, since in low-attention states mice might commit to a difficulty-independent heuristic strategy.

The analysis of angles between state axes highlighted a large angular separation between variables, with sensory and movement axes having the smallest separation. This latter observation agreed with previous reports both at the local scale of small neuronal assemblies [50], and at the mesoscale [35], indicating a covariability axis between these variables. Similarly, the smaller angles observed between the movement axis and the attentional axis agreed with a recent report showing that attention enhances distinctive spatial features in movement-related activations across these cortical regions [35].

### RNN implementation and mechanistic insights

Recurrent state-space models, including RNNs, have been previously used in mechanistic investigations of decisionmaking processes [23, 24]. Moreover, representational similarity analysis of the state-space of RNNs and neural responses has been successfully used to infer underlying computations [23]. Here, we adopted a similar approach, but with three main distinguishing features. First, we trained the network with the animals’ decisions, rather than the task rule. This constituted a significant departure from previous research, which added noise to fully-deterministic RNNs to capture logistic behavioral tuning functions [24, 51]. Instead, we trained the network with “contradictory” information, such as that involved in the inconsistent trial-to-trial animals’ decisions, thus exposing the network to the biases and heuristics of the animals. Training with the animal choices was akin to training with label noise, for which many deep learning algorithms are robust [52]. The RNN outputs effectively implemented two dynamic accumulators providing time-dependent scores for L/R choices, with the difference between the scores being proportional to the psychometric function. This result was probably related to the mathematical observation that if L/R choices were determined by two accumulators (for the left or right evidence, respectively), the log-likelihood ratio of the conditional probabilities for a specific choice, given the state of the accumulators, can be shown to be proportional to the psychometric (logistic) function [53, 54]. The second novelty was that we trained the RNN to learn an invariance regarding absolute orientations, which were uninformative for the task choice. Finally, the third novelty concerned attentional modulations. As observed in the neural data, the added attentional input to the RNN caused an isomorphic shift of the decision-making manifold, which retained the geometry of the decision variable. Geometry-preserving isomorphic shifts in low dimensional embedding spaces, might reflect a general ‘decorrelation’ principle for variables that are concurrently represented across overlapping cortical networks. These results confirmed that, mechanistically, the representational dynamic of choice signals reflected the decision-making computations underlying the animals’ psychophysical behavior.

### Limitations and open questions

Our results raise several questions to be addressed in future studies; for instance, it is unclear whether the broad distribution of choice signals mirrored equally broad spiking activations. Regarding anatomical considerations, feedback signals are known to preferentially target deep layers (five and six) and layer one [55].Considering that our imaging macroscope focused on superficial cortical layers and that GCaMP was expressed across the cortex, it is possible that choice signals reflected long-range axon-terminal activations and /or depolarizations in apical dendritic trees, rather than somatic firing. Concurrent imaging and electrophysiological recordings across layers would clarify this point.

Our study relied on correlative measures, therefore loss- and gain-of-function perturbative experiments will be necessary to establish causality. Of particular interest would be the inactivation of lateral visual areas in view of the observed ventral-stream prominence. Furthermore, patterned optogenetic methodologies with single-cell resolution might enable the stimulation of those neurons that most significantly carry choice-relevant information in these regions.

In sum, broadly distributed feedback decision signals, having a representational dynamics consistent with decisionmaking computations underlying the perceptual task, might represent an activity—and computational—substrate capable of “molding” early sensory processing and sensory to decision transformations depending on the underlying decisionmaking process for probabilistic-inference computations in changeable agent-environment interactions [56].

## ACKNOWLEDGEMENTS

We thank Rie Nishiyama, Yuka Iwamoto, and Yuki Goya for technical support in multiple aspects of the experiments. We thank O’Hara and CO., LTD., for their support with the equipment. This work was funded by RIKEN BSI and RIKEN CBS institutional funding; HFSP postdoctoral fellowship LT000582/2019 to J.O.; JSPS grants 26290011, 17H06037, 372 C0219129 to A.B.; and a Fujitsu collaborative grant.

## Author Contributions

A.B. and J.O. conceived the project and wrote the manuscript. M.A., R.A. and D.L. collected the data. M.A. and J.O. pre-processed the data and J.O. analyzed the data.

## Competing Interests

The authors declare no competing interests.

## Materials and methods

### Experimental procedures

Details of the experimental procedures (surgeries, behavioral training, recordings of body and eye movements, imaging methods, and preprocessing of fluorescence data) have been described in Ref. [35]. We summarize them here in brief.

### Surgeries

All surgical and experimental procedures were approved by the Support Unit for Animal Resources Development of RIKEN CBS. Transgenic mice used in this work were Thy1-GCaMP6f mice *(n* = 7). For all reported results, the number of sessions per animal ranged from 9 to 60, with a minimum and maximum number of trials per animal from 1000 to 8000. Animals were implanted with a cranial post for head fixation and a round chamber (~6 mm diameter) for optical access to neural recordings.

### Behavioral training

Animals were trained in a 2AFC orientation discrimination task. Two oriented Gabor patches were shown on the left and right side of a screen positioned in front of the animal at ±35° eccentricity relative to the body’s midline. Mice had to report which of the two stimuli matched a target orientation (vertical, *n* = 5; horizontal, *n* = 2). The smallest orientation difference varied depending on animals, from 3° to 30°. The largest difference—the easiest discriminations—was ±90°. Animals signaled their choice by rotating a rubber wheel with their front paws, which shifted stimuli horizontally on the screen. For a response to be correct, the target stimulus had to be shifted to the center of the screen, upon which the animal was rewarded with 4 μl of water. Incorrect responses were discouraged with a prolonged (10 s) inter-trial interval and a flickering checkerboard stimulus (2 Hz). If no response was made within 10 s (time-out trials), neither reward nor discouragement was given. Animals were imaged after exceeding a performance threshold of 75 % correct rate for 5-10 consecutive sessions. To work with a coherent behavioral dataset, we excluded sessions with exceedingly large fractions of time-outs (≥20 %) or with average performance dropping below 60 %. Every trial consisted of an open-loop period (OL: 1.5 s) and a closed-loop period (CL: 0 s to 10 s), followed by an inter-trial interval (ITI: 3 s to 5 s randomized). We recorded cortical responses, wheel rotations, and eye/pupil videos from a pre-stimulus period (1 s duration) until the end of the trial. Stimuli were presented in the OL period, when wheel rotations did not produce any stimulus movement.

### Saccades, pupil area, and body movements

We monitored the contralateral eye using a CMOS camera. Automatic tracking of the pupil position was done with custom software (Matlab, Mathworks). We confirmed the accuracy of pupil tracking by visually inspecting hundreds of trials. Saccade detection was achieved by applying an adaptive elliptic thresholding algorithm to saccade velocities. We discarded the saccades that lasted ≤60 ms and were smaller than 1.5°. We extracted the time, magnitude, duration, velocity, start, and landing positions of each saccade. We calculated the average pupil area for each imaging session by averaging area values across all trials within the session. Pupil area in every trial was z-scored for each session.

### Wheel detection

We recorded wheel rotations with a rotary encoder and flagged as potential wheel movement time points when the velocity had a zero-crossing (i.e., sign change) and deviated from zero above a fixed threshold (20°).

### Imaging

Mice were placed under a macroscope for wide-field imaging in tandem-lens epifluorescence configuration. We imaged GCaMP6f fluorescent signals using a CMOS camera (pco.edge 5.5).

### Pre-processing of fluorescence data

We motion corrected GCaMP data using a semi-automated control-point selection method (MATLAB cpselect). To compute relative fluorescence responses, we calculated a grand-average scalar 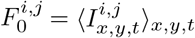, with 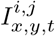 the XYT image tensor in trial *i*, session *j*. We then used this scalar to normalize the raw data tensor 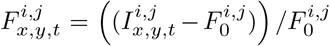. The data in each trial were then band-pass filtered (0.1 Hz to 12 Hz). Each tensor was compressed with spatial binning (130 μm × 130 μm) with 50 % overlap). Trial data was further downsampled to 30 fps and low-pass filtered with a cutoff at 8 Hz.

### Retinotopy

We used a standard frequency-based method [57] with slowly moving horizontal and vertical flickering bars. Visual area segmentation was done based on azimuth and elevation gradient inversions. To center and orient maps across animals, we used the centroid of area V1 and the iso-azimuth line passing through it.

### Alignment to the Allen Mouse Brain Common Coordinate Framework

Imaging data from each animal was aligned to the Allen Mouse Brain Common Coordinate Framework (CCF) following the approach described by Waters [58] In brief, we extracted the centroids of areas V1, RL and PM, using them to create a triangle that we aligned to the one from the Allen CCF by first making the V1 vertices coincide. We then rotated and scaled the triangle to minimize the distance between the other vertices while maintaining the original angles.

### Data Analysis

#### LocaNMF decomposition

LocaNMF analysis was done following the methods described by Saxena [21]. In brief, imaging data cross all trials and sessions was first concatenated and its dimensionality reduced using singular-value decomposition (SVD) up to 99 % of the original variance. LocaNMF was initialized using 10 regions based on the Allen CCF, and corresponded to: V1 (VISp), PM (VISpm), AM (VISam), A (VISa), SSt (SSp-tr), RL (VISrl), SSb (SSp-bfd), AL (VISal), VL (VISl and VISli), RS (RSPagl and RSPd) (Extended Data Fig. 1). A mask in each area was created by setting a distance *D* = 1 within the region boundaries and an exponential decrease (to zero) for pixels outside the boundary. The localization penalty for each pixel was 1 – *D* (Extended Data Fig. 1). LocaNMF Rank Line Search was run with these 10 regions with a localization threshold of 75 %, and total explained variance of 99 %, resulting on average in approximately 200 components per animal. After decomposition, temporal components were split back into the original trial structure. More formally, LocaNMF produces a decomposition tensor 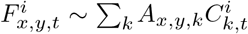 for trial *i*. Where *A_x,y,k_* is the spatial part of component *k*, and 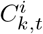 it’s temporal part. Each spatial part of the components is significantly localized and it can be mapped onto its original seeding area. The temporal component captures the unique trial to trial variability, and all subsequent analysis in the time domain have been done using only the temporal part 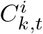 of locaNMF components. Total explained variance (EV) of the decomposition, was computed by projecting one component at a time across the whole time series and checking the relative explained variance across pixels with the original recording.

#### Global State Vector definition

We defined a state vector as a one-dimensional projection of locaNMF temporal components **C**(*t*) that maximizes the weighted distance between the trajectories of two trial groups *A* and *B* (herein after bold letters indicate vectors). For each trial group, we defined average trajectories 〈**A**(*t*)〉 and 〈**B**(*t*)〉 and define 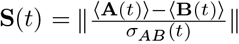 where *σ_AB_*(*t*) is the pooled standard deviation between the two groups. State vector projections for the *i*-th trial were then obtained by the dot product *P^i^*(*t*) = **S**(*t*) · **C**^*i*^(*t*), where **C**^*i*^(*t*) are the temporal locaNMF components of trial *i*. Discriminability between the original A and B groups is then computed as 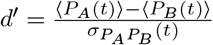 i.e., the difference between the averaged projections of groups **A** and **B**, divided by their pooled standard deviation. To validate state vectors, projections, and discriminability, we performed 5-fold cross validation, i.e., state vectors were defined using only 20 % of the trials of each group and projections and *d!* were computed on the remaining trials.

#### State-vector stability

To assess the stability of state vectors, we computed instantaneous state vectors **S**(*t*) using a ‘backward’ 3-frame averaging window (~100 ms) and then computed its autocorrelation *C*(**S**(*t*), **S**(*t*′)). For sensory, movement, and sustained attention state vectors we chose time-independent state vector **S** = **S**(*t*\*), where *t*\* was chosen within the largest stability cluster (represented by a gray bar in the respective figures). For the state vector of choice we used the original **S**(*t*) to keep track of when choice information first appeared and whether its signature was unique.

#### Area-specific state vectors

We defined state vectors for each of the 10 areas by only using the subset of locaNMF components 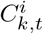 that originated from that area. This is akin to first projecting onto the subspace defined by the components of a particular area, and then obtaining the associated state vector.

#### Task-related state vectors

**Stimulus:** For stimulus statevector we used for the first group *A* all the trials in the time interval −0.5s to 0.5s centered on stimulus onset. For group *B*, since all trials had stimulus presentation, we took the same trials on the – 1 s to 0s interval before stimulus presentation. **Contralateral stimulus:** We used as group *A* the trials where the left stimulus was horizontal and as group *B* the trials where the left stimulus was vertical (Extended Data Fig. 3). **Wheel movement:** Group *A* consisted of trials whose first movement after stimulus presentation occurred at least 0.5 s after stimulus onset and without any saccades detected in the previous 0.5 s. These trials were aligned to the detected movement onset. Group *B* consisted of trials with no movement detected on the first 5 s after stimulus onset. These trials were aligned by taking a random frame within that same interval. **Saccades:** Defined as for wheel movements. **Sustained attention:** Sustained attention was measured by changes in pupil area during the trial. We computed the pupil change measure, *pA*, as the difference between the maximum pupil area after stimulus onset and the average area 1 s before stimulus onset. We labeled ‘high attention’ trials (group *A*) those in the top 33-th percentile of the *pA* distribution and ‘low attention’ (group *B*) those in the bottom 33-th percentile. **Choice:** L/R choices in each trial were measured from the direction of the first movement after stimulus onset. The state vector for group *A* was computed from right-choice trials and for group *B* using left-choice trials. As for the detection of wheel movements, we restricted the analysis to trials where the first movement happened at least 0.5 s after stimulus onset and with no saccades 0.5 s before the detected movement. Trials were aligned to the time of movement detection.

#### Piecewise linear fitting of d′ curves

To fit the time-evolving *d*′ curves on periods before and after movement onset, we performed a 2-slope piecewise linear fitting using the Shape Language Modeling toolbox (MATLAB Central File Exchange, John D’Errico (2021). SLM - Shape Language Modeling). The method performs two linear fits in a fixed interval with a single knot between them. We chose the interval – 1 s before movement onset up to 95-th percentile of the peak post-movement response amplitude (typically occurring around 0.3 s after movement onset). The position of the knot determined the slope change time.

#### Spatial-Distribution Index (SDI)

The SDI for a given state vector was computed as 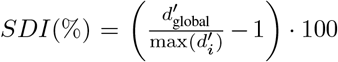 where 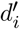 refers to the area-specific *d*′ scores.

### Recurrent Neural Network model

The Recurrent Neural Network (RNN) consisted of a single RNN module with *N* =128 neurons (ReLU activations), receiving 3 inputs (left stimulus, right stimulus, attention level) and producing as an output a binary response for left or right choices (softmax activation).

#### Inputs

The input space consisted of a sequence of 25 frames. Stimulus orientations were mapped to the range (−1 +1) corresponding to −90° to 90° and were presented after the first 10 frames. The difficulty of a trial was encoded by the absolute difference between the two stimulus signals. Attention was modeled as a constant binary signal (0 or 1), already present at the beginning of the trial. A small noise (normally distributed with amplitude 0.01) was added to the input signals to improve the robustness of the optimization, but it was irrelevant for the psychometric fits.

#### Training

For training the network we generated simulated animal responses by computing left or right choices following a psychometric curve of the form 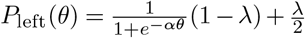 where *θ* is the difference between the 2 inputs, *λ* the lapse rate, and *α* controls the slope. We used a constant *λ* = 0.1 and *α* = 3/90 for low attention and *α* = 10/90 for high attention. That is, attention decreased the amount of label noise during training. To train the network we used 6400 trials per difficulty level and chose 13 difficulty levels with angle differences uniformly distributed from −90° to 90°. We trained the network using a batch normalization layer and a custom loss function consisting of the categorical cross entropy at the time of stimulus presentation and at the last frame. We included the stimulus presentation time in the loss function to force the output to not drift before stimulus presentation, following a previous procedure [23]. Accuracy during training was computed by the categorical accuracy at the end of the trial. The network was implemented with TensorFlow 2.0 and trained using the Adam optimizer for 25 epochs with a batch size of 640. Note that training the network with the animal choices made the network robust to overfitting. We trained 10 different networks by generating a new set of inputs and randomly initializing the networks weights.

#### Analysis

We analyzed the output of the RNN in the same way we did with the neural data, but using the time series of the *N* = 128 neurons instead of the locaNMF components to define choice and attention state vectors.

---

**Extended Data Fig. 1.**
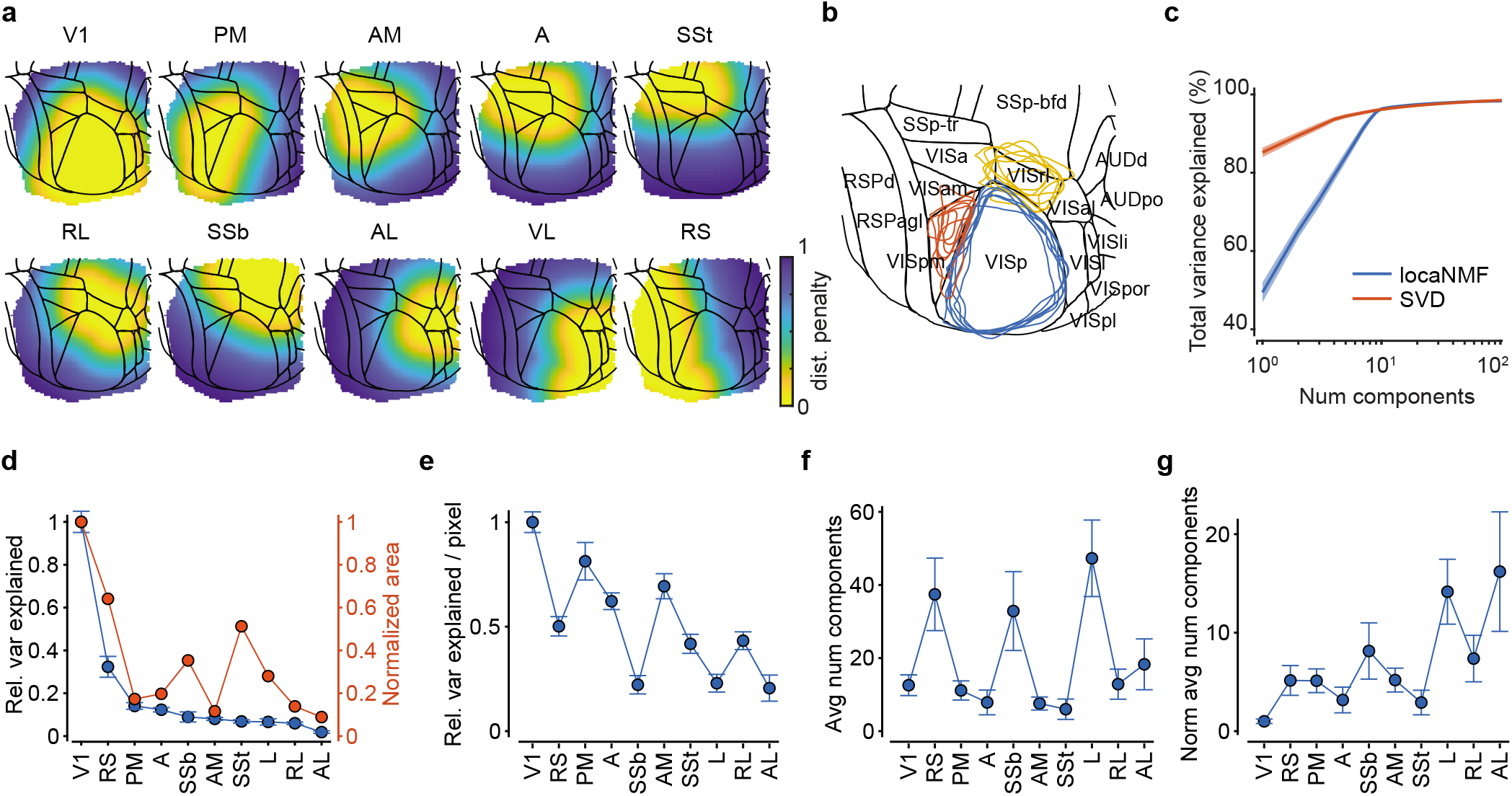
Statistics of locaNMF decomposition. **a**, Pixel-wise distance penalty maps used to initialize the ten regions for the locaNMF decomposition. Penalties within the boundaries of each region were 0 and increased exponentially with distance from the boundary. **b**, Superposition of our retinotopically aligned animals (blue, red, yellow contours) with the Allen Brain Atlas. **c**, Total variance explained as a function of the number of components for locaNMF and standard singular-value decomposition (SVD) for a given animal. **d**, Blue = relative variance explained with respect to V1 for each of the areas ordered by their variance explained. Red = surface areas relative to V1. **e**, Fraction of the total variance explained by the locaNMF components from each region normalized by the number of pixels in each region. **f**, Number of components required in each area to reach 99 % of total explained variance, averaged across animals. This number does not simply reflect area sizes. For instance, VL decomposition resulted in a large number of components in all animals despite being a small region. **g**, Average number of components as in **f**, normalized by the size of each region.

**Extended Data Fig. 2.**
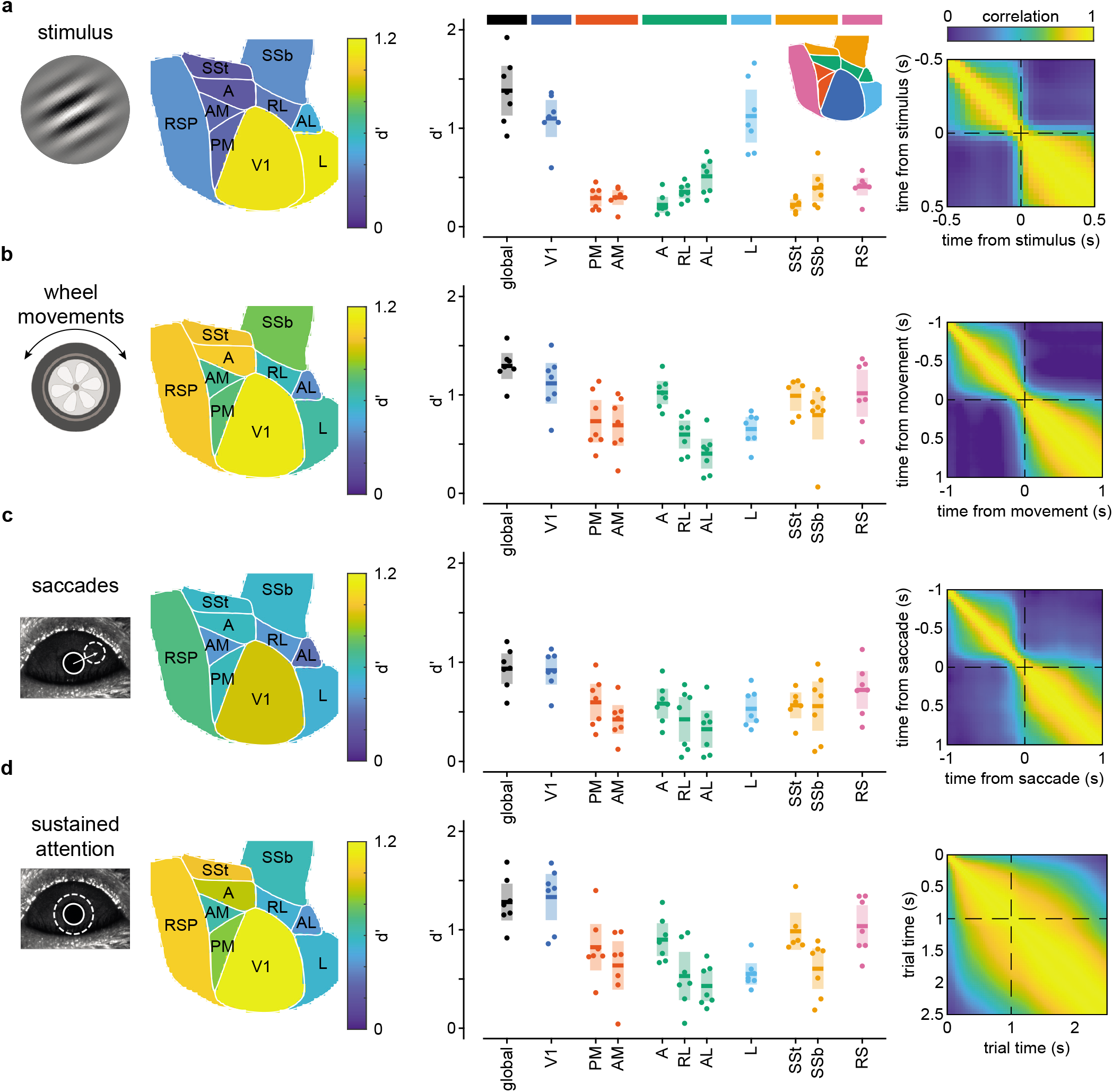
Sensory, movements and top-down discriminability statistics across areas. **a**, Average discriminability *d*′ of each area for stimulus, wheel movements, saccades, and sustained attention (top to bottom rows). **b**, Statistics of global *d*′ values (ignoring source location), and for each individual area across animals. Individual dots for each animal, middle bar mean, and shaded area 95% CI of the mean. Inset: color-code reference for each of the areas. **c**, Stability of each global state vector in time for a representative example animal. For stimulus, wheel movements, and saccades, the state vector becomes stable right after the event onset, whereas for sustained attention it is stable throughout the trial.

**Extended Data Fig. 3.**
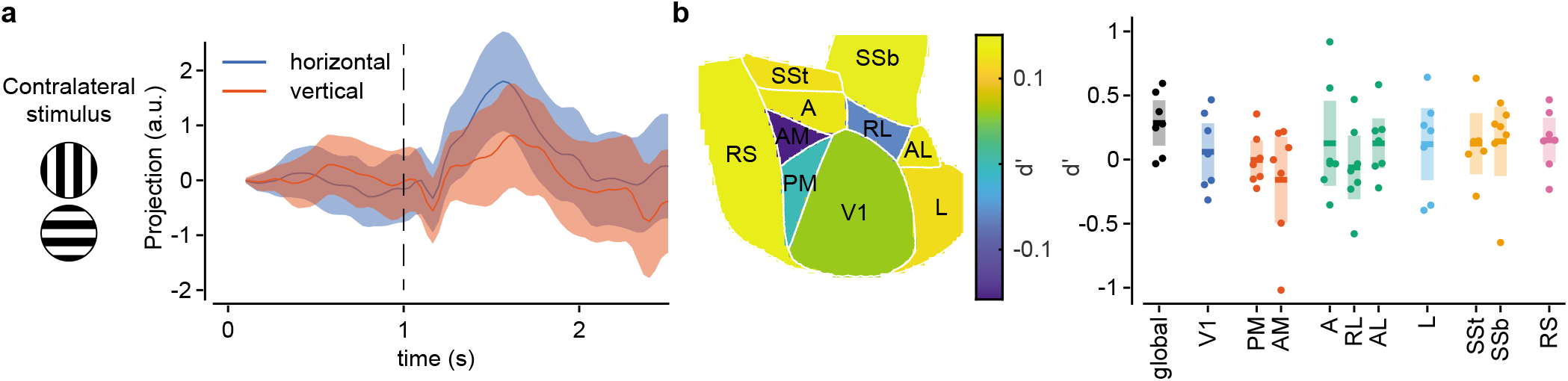
Widefield activity contained no information about the contralateral stimulus orientation. **a**, Time dependence of response projections onto a state vector defined using horizontal or vertical contralateral stimulus trials for a representative animal. Trajectories did not split throughout the trial. Line is the trial average and shaded areas s.e. **b**, Statistics of global and areaspecific *d*′ for the same state-vector. All area-based *d*′ values were consistent with no discriminability power; each dot is one animal, middle bar and shaded area are the average across animals and 95 % CI.

**Extended Data Fig. 4.**
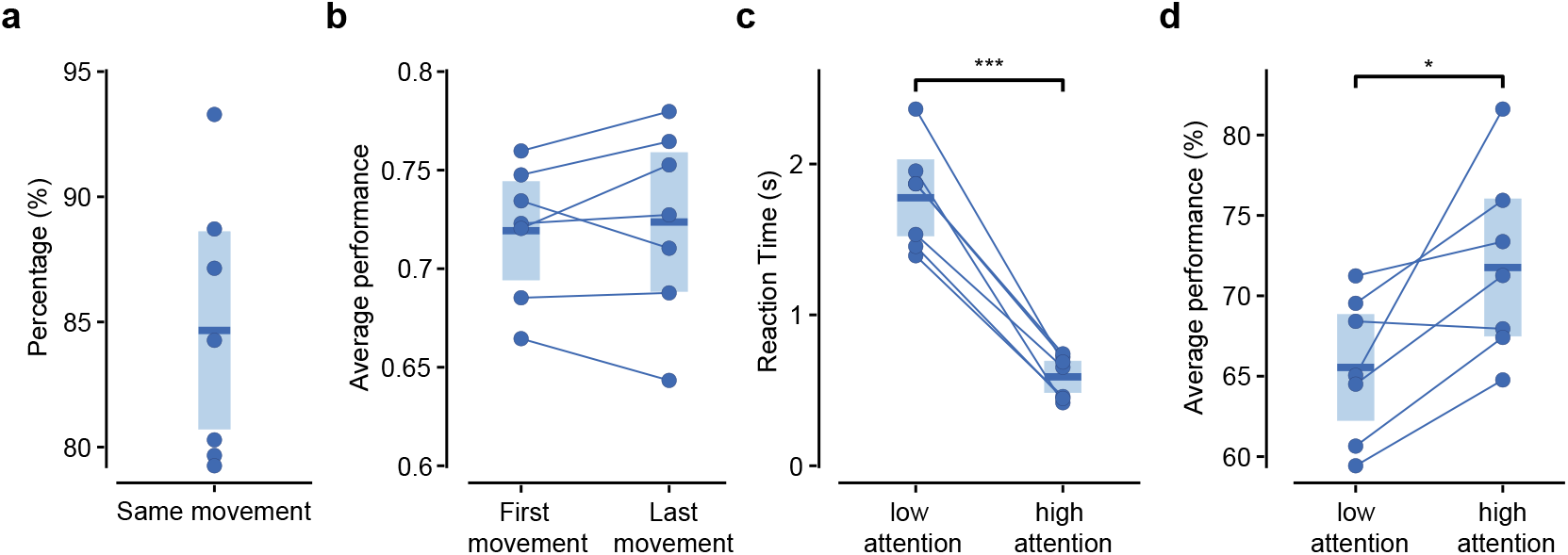
First-movement correlation with choice and attention-dependent performance. **a**, Fraction of occurrences when the direction of the first wheel movement coincided with the direction of the last movement in the trial (i.e., the movement that ended the trial). **b**, Comparison of overall performance when considering either the first or the last movement (Paired t-test, *p* = 0.8). **c**, Reaction times for the first movement depended on attention, being significantly shorter in high attention trials (paired t-test, *p* = 4 · 10 – 5). **d**, Average performance was consistently higher in high-attention trials (paired t-test, *p* = 0.02). In all panels, each dot indicates one animal; middle bar and shaded area are the average across animals and 95 % CI.

**Extended Data Fig. 5.**
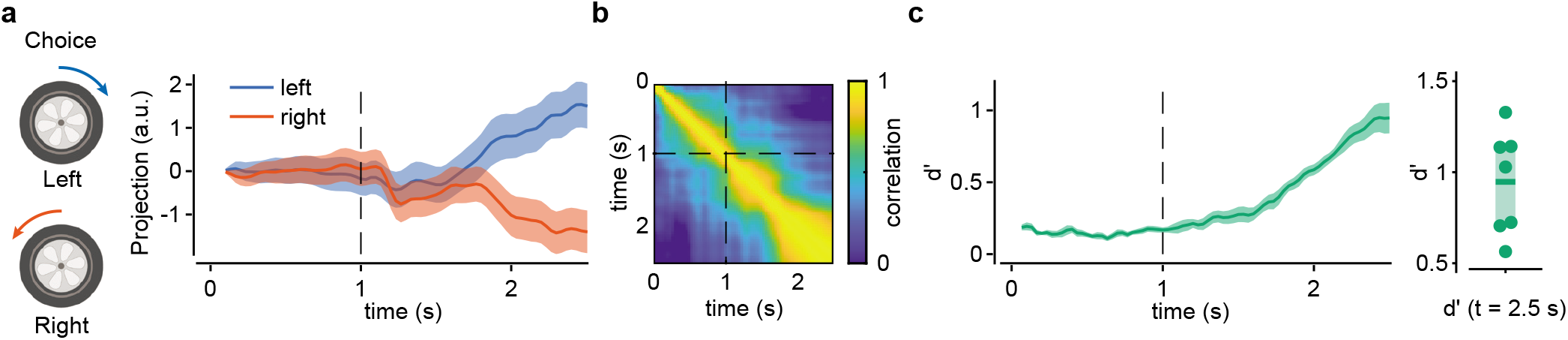
Choice signals can be distinguished in trial time. **a**, Projections onto the state vector for choice defined in trial-time—instead of movement time— for a representative animal. Only trials where the first movement appeared in the 1.5 s to 2.5 s window were included. Trajectories started to split within the same window **b**, Stability of the choice state vector in trial time. A first signature of stability appears soon after stimulus onset. **c**, Global *d*′ evolution for the same state vector averaged across animals (left), and statistics of peak values (right; each dot is one animal, middle bar and shaded area are the average across animals and 95 % CI). *d*′ starts to increase right after stimulus onset and before movement onset.

**Extended Data Fig. 6.**
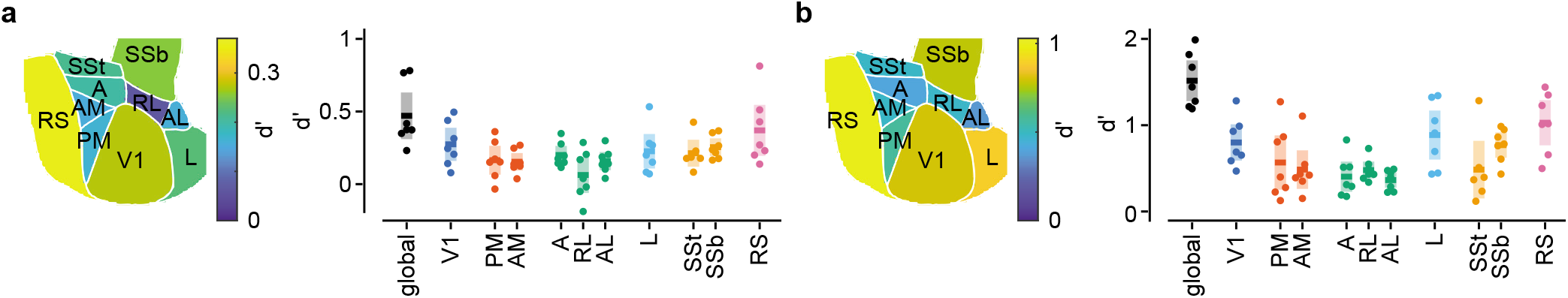
Choice-specific area contributions. **a**, and **b**, Global and area-based *d*′ statistics for the choice state vector in movement time (Fig. 3), before movement onset (left, t = −0.1 s), and peak values after movement onset (right, t=0.3s).

**Extended Data Fig. 7.**
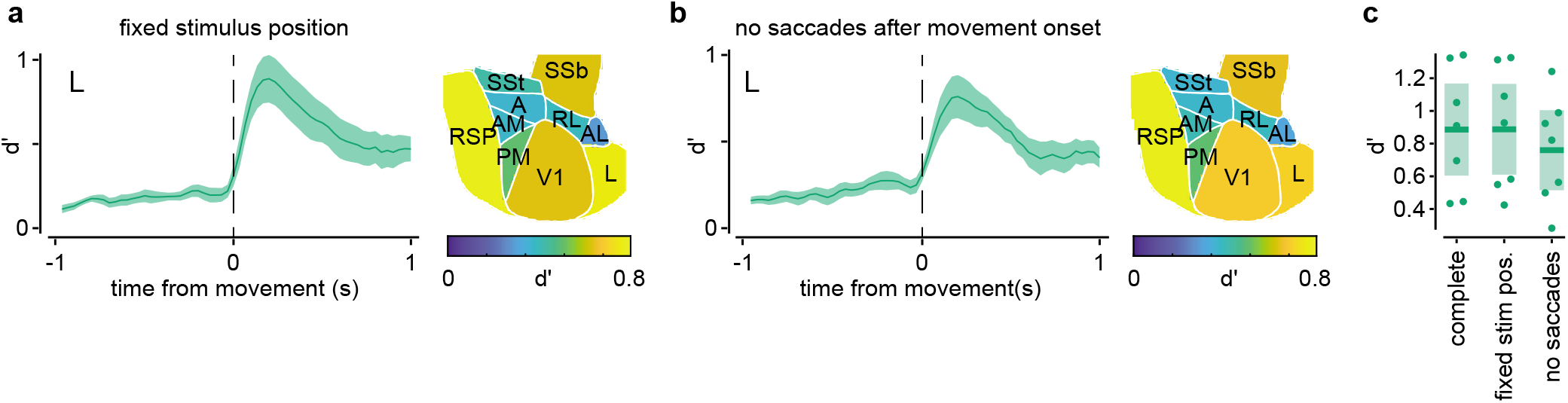
Ventral stream choice signature is not linked to eye or stimulus movements. **a**, left: Evolution of choice *d*′ for area L during trials where the first movement happened within 1s of the stimulus presentation, i.e., the stimulus was always on the same position in the screen. Right: area-specific *d*′ 0.2 s after movement for the same trials. **b**, Same as in **a**, but also with the constraint that there were no saccades 0.5 s before or after the movement onset. **c**, Comparison of peak *d*′ values in area L for the three controlled conditions: complete (same as Fig. 3 on the main text), and those shown here in panels **a** and **b**.

## Notes

### Competing Interest Statement

The authors have declared no competing interest.

